# Stepwise bladder carcinogenesis reveals transcriptional reprogramming involving MMP1 and COL7A1

**DOI:** 10.64898/2026.09.04.749539

**Authors:** Yuko Nagashima, Haru Yamamoto, Mohamed Elbadawy, Yoshiko Naito, Amira Abugomaa, Ryouichi Tsunedomi, Masahiro Kaneda, Tatsuya Usui, Kazuaki Sasaki

## Abstract

Bladder cancer develops through multistep molecular alterations, yet the events underlying tumor initiation and progression remain poorly understood. Here, we reconstructed stepwise bladder carcinogenesis using canine bladder organoids combined with chemical carcinogenesis and serial xenotransplantation. Carcinogen-exposed normal organoids generated benign tumors that subsequently progressed to invasive carcinomas. Despite a low tumor mutational burden and limited acquisition of cancer-associated mutations, malignant progression was accompanied by extensive transcriptional reprogramming, including enrichment of epithelial-to-mesenchymal transition programs. COL7A1 and MMP1 were progressively upregulated during progression, and knockdown of either gene suppressed organoid proliferation and invasion. These functional dependencies were reproduced in independently established spontaneous canine bladder cancer organoids, where silencing of either gene also reduced xenograft growth and mitosis-related gene programs. MMP1 knockdown decreased COL7A1 expression, suggesting a potential regulatory relationship. MMP1 was also elevated in basal/squamous human bladder cancers, and its depletion suppressed human bladder cancer cell proliferation. These findings identify transcriptional reprogramming and MMP1/COL7A1 dependencies as prominent features of malignant progression.

## Introduction

Bladder cancer is one of the major malignancies in both humans and dogs^1–3^, and muscle-invasive bladder cancer (MIBC) is a highly aggressive disease of substantial clinical importance^4,5^. Canine urothelial carcinoma shares many features with human MIBC, including clinical behavior, histopathological characteristics, and treatment response^6–10^. Naturally-occurring tumors in companion animals arise in a shared environment with humans and are regarded as important translational models for cancer research. This concept underlies comparative oncology^6,11–14^. At the molecular level, canine urothelial carcinoma exhibits similarities to human bladder cancer^8,15,16^, while also harboring canine-specific features, such as a high frequency of BRAF mutations^17–20^.

Bladder cancer is a malignancy influenced by exogenous factors, including exposure to chemical substances such as aromatic amines derived from smoking^21–23^, occupational exposure^21,22,24^, and environmental pollution^22,25–28^. Its initiation and progression are thought to be driven by the accumulation of multistep molecular alterations^29–31^. During tumor progression, phenotypic changes such as epithelial–mesenchymal transition (EMT) and extracellular matrix (ECM) remodeling play important roles^32–35^. However, the transition from normal urothelium to tumor formation and the molecular mechanisms underlying early carcinogenesis remain incompletely understood^36–38^.

Bladder cancer research has traditionally relied on cultured cell lines and animal models. However, cultured cell lines are patient-specific, subject to selection pressure during long-term passaging^39–41^, and do not necessarily reflect tumor heterogeneity or tumor development^42,43^. Animal models, while capable of recapitulating the *in vivo* environment, are limited in their ability to analyze molecular changes during tumor progression in a time-resolved manner^44–46^. Chemical carcinogenesis has long been used as a principal experimental approach in cancer research, and long-term *in vivo* carcinogenicity assays are established methods for evaluating tumor development^47–50^. However, these models require extended periods^51^ and make it difficult to investigate the molecular mechanisms underlying carcinogenesis, particularly during the early stages of tumor development^52–55^. Therefore, experimental models are needed that enable stepwise analysis of tumor development to better understand carcinogenesis and its molecular mechanisms^56–58^.

Organoids have become established experimental models for investigating cancer biology, disease mechanisms, and therapeutic responses^59–61^. Patient-derived bladder cancer organoids retain the histological features, genetic alterations, and drug responsiveness of the original tumors and are used for the analysis of cancer biology and therapeutic responses^62–64^. In our previous studies, we established organoid models from several spontaneous canine cancers, including bladder cancer, prostate cancer, lung cancer, mesothelioma, and apocrine gland anal sac adenocarcinoma, and demonstrated that these models faithfully recapitulate key features of the original tumors and provide valuable platforms for translational cancer research^65–69^. However, most organoid models are established from pre-existing tumors and therefore do not recapitulate the stepwise process of carcinogenesis from normal epithelium to tumor formation^58^. In addition, organoids have limitations in modeling molecular pathways that depend on the tumor microenvironment, such as angiogenesis and immune responses, and may not fully reflect cancer progression^55,58,61^. Organoid-based carcinogenesis models induced by chemical agents have also been reported^70–73^, but such approaches remain limited in bladder epithelium, and models that recapitulate stepwise progression from normal urothelium to tumor formation and malignancy are still lacking^58^.

We previously established urine-derived normal canine bladder organoids from healthy beagle dogs and demonstrated their utility as a physiologically relevant *in vitro* model of the urothelium.^74^ However, whether these organoids could be used to reconstruct the multistep process of bladder carcinogenesis and thereby elucidate the molecular changes accompanying malignant progression remained unclear. Here, we reconstructed stepwise bladder carcinogenesis by combining chemical carcinogen exposure of normal canine bladder organoids with serial xenotransplantation. Using this model, we longitudinally characterized genomic and transcriptional changes during malignant progression and found that extensive transcriptional reprogramming, including enrichment of epithelial-to-mesenchymal transition (EMT), occurred despite limited acquisition of cancer-associated genomic alterations. We further identified MMP1 and COL7A1 as functional mediators of malignant phenotypes and validated their relevance in independently established spontaneous canine bladder cancer organoids and human bladder cancer.

## Materials and methods

### Generation of CNBO

Canine normal bladder organoids (CNBO) were established from urine-derived cells collected via catheterization from a clinically healthy female beagle dog (7 years old, 9 kg), as previously reported^74^. Urine samples were centrifuged at 600 × g for 3 min at room temperature (RT). Cell pellets were washed with PBS and embedded in 40 μL of Matrigel (Corning, Corning, NY, USA) at 8 × 10^3^ cells per well in 24-well plates. Cells were cultured in organoid growth medium as previously described^74^. For passaging (every 7–10 days), Matrigel was dissolved with 5 mM EDTA in PBS on ice for 1.5 h. Organoids were dissociated into single cells using TrypLE Express (Thermo Fisher Scientific, Waltham, MA, USA), neutralized with fetal bovine serum (FBS; Thermo Fisher Scientific), filtered through a 70-μm cell strainer, counted, and reseeded.

### Hematoxylin and eosin (H&E) staining

Organoids and tumor tissues were fixed in 4% paraformaldehyde at RT for 24 h, followed by dehydration through a graded ethanol series and xylene, and subsequent paraffin embedding. Sections (4 μm thick) were prepared and stained with hematoxylin and eosin according to standard protocols. Images were acquired using a light microscope (BX-52; Olympus, Tokyo, Japan).

### Immunohistochemical staining

Immunohistochemical (IHC) staining was performed as previously reported^69^. Antigen retrieval was performed using 10 mM citrate buffer (pH 6.0) heated at 121 °C for 5 min. After cooling, endogenous peroxidase activity was blocked with 1% hydrogen peroxide for 30 min at RT. Sections were blocked with 10% normal goat serum for 30 min and incubated with primary antibodies (listed in Table 1) at 4 °C overnight. After washing with PBS, sections were incubated with secondary antibodies, and immunoreactivity was visualized using a diaminobenzidine (DAB) substrate (Nacalai Tesque, Kyoto, Japan). Sections were counterstained with hematoxylin, and images were captured using a microscope (BX-52).

**Table 1.**
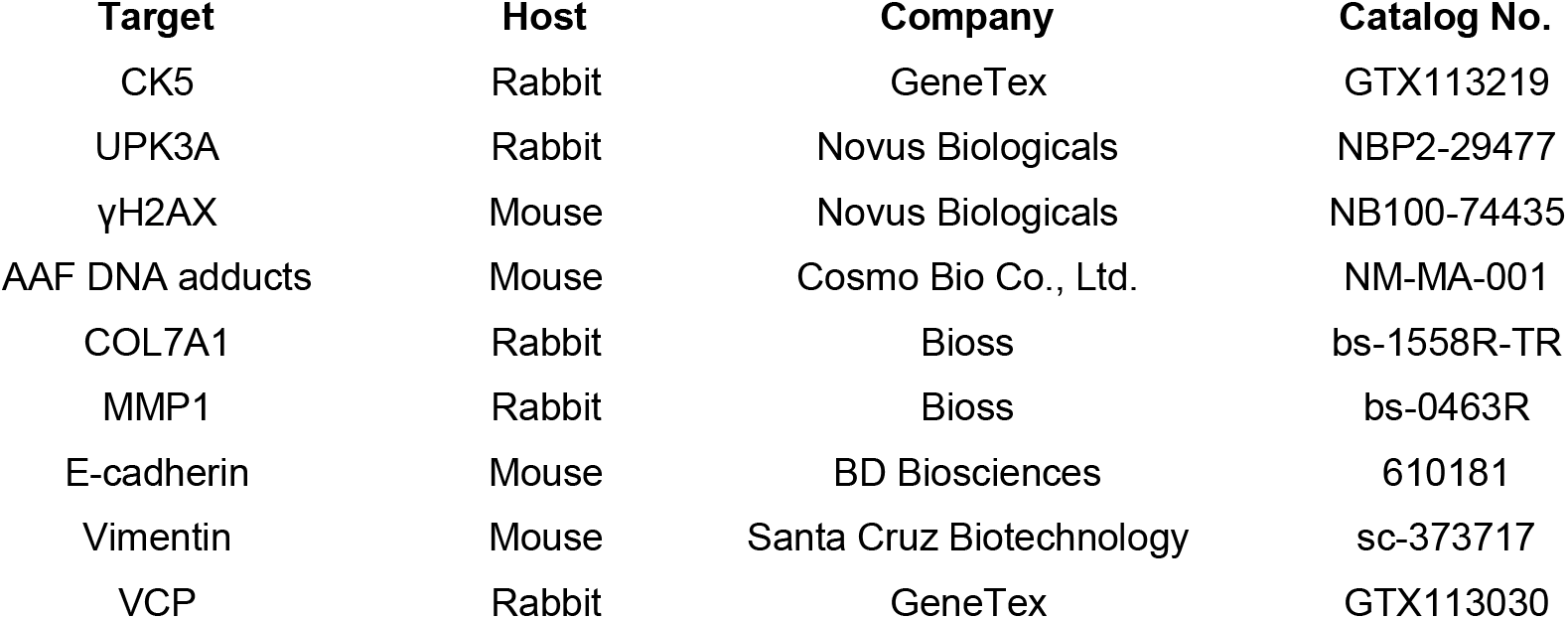
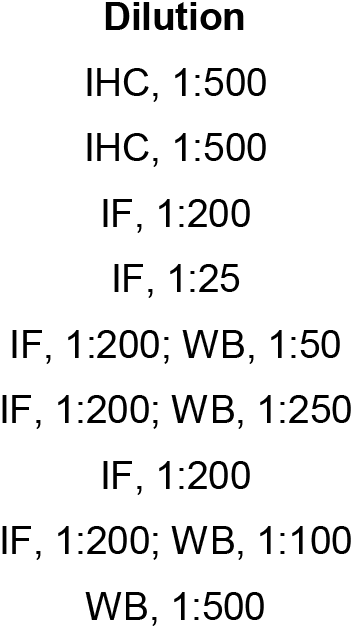
Primary antibodies used in this study.

### Chemical treatment of CNBO with 2-AAF

CNBOs were seeded at 8 × 10³ cells per well in 24-well plates and allowed to establish for 24 h before treatment. CNBOs embedded in Matrigel were treated with 2-acetylaminofluorene (2-AAF; Tokyo Chemical Industry, Tokyo, Japan) dissolved in dimethyl sulfoxide (DMSO; FUJIFILM Wako, Osaka, Japan). To determine an appropriate treatment concentration, dose–response experiments were conducted using 0.01, 0.1, 1, and 10 μM 2-AAF. For cell viability assays, CNBOs were dissociated into single-cell suspensions, embedded in Matrigel, and seeded into 96-well plates at a density of 1 × 10³ cells per well in triplicate. After 24 h of culture, cells were treated with the indicated concentrations of 2-AAF. Culture medium containing the corresponding concentration of 2-AAF was replaced at 72 h. After 144 h of treatment, cell viability was assessed using a PrestoBlue Kit (Thermo Fisher Scientific) according to the manufacturer’s instructions. Fluorescence intensity was measured using a microplate reader (Tecan Group Ltd., Männedorf, Switzerland), normalized to vehicle-treated controls, and used to estimate the half-maximal inhibitory concentration (IC₅₀). Based on the results of the dose–response experiments, 1 μM 2-AAF was selected for subsequent experiments. Vehicle control organoids were treated with an equivalent concentration of DMSO.

### Quantitative real-time polymerase chain reaction

Total RNA was extracted from cells using the FavorPrep Tissue Total RNA Purification Mini Kit (Favorgen Biotech Corp., Pingtung, Taiwan) following the manufacturer’s protocol. First-strand cDNA was synthesized using the ReverTra Ace qPCR RT Kit (TOYOBO, Osaka, Japan). Quantitative real-time PCR was performed using the QuantiTect SYBR Green PCR Kit (QIAGEN) on a StepOnePlus Real-Time PCR System (Applied Biosystems, Waltham, MA, USA). Relative gene expression levels were calculated using the 2^−ΔΔCt method. Primer sequences are listed in Table 2.

**Table 2.**
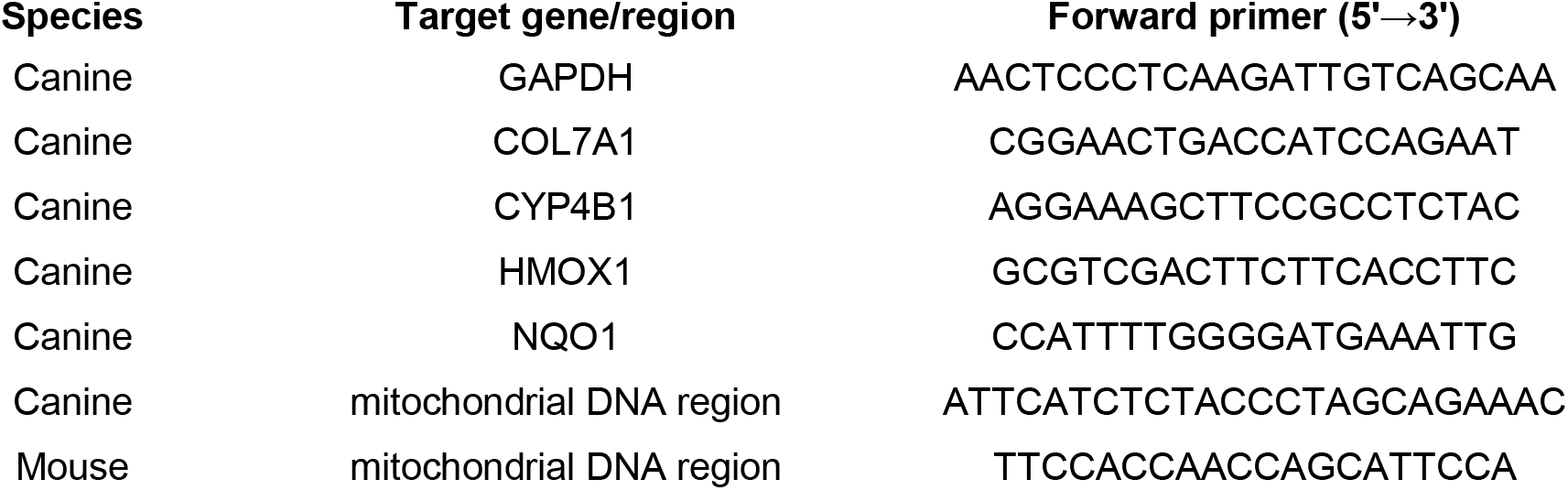
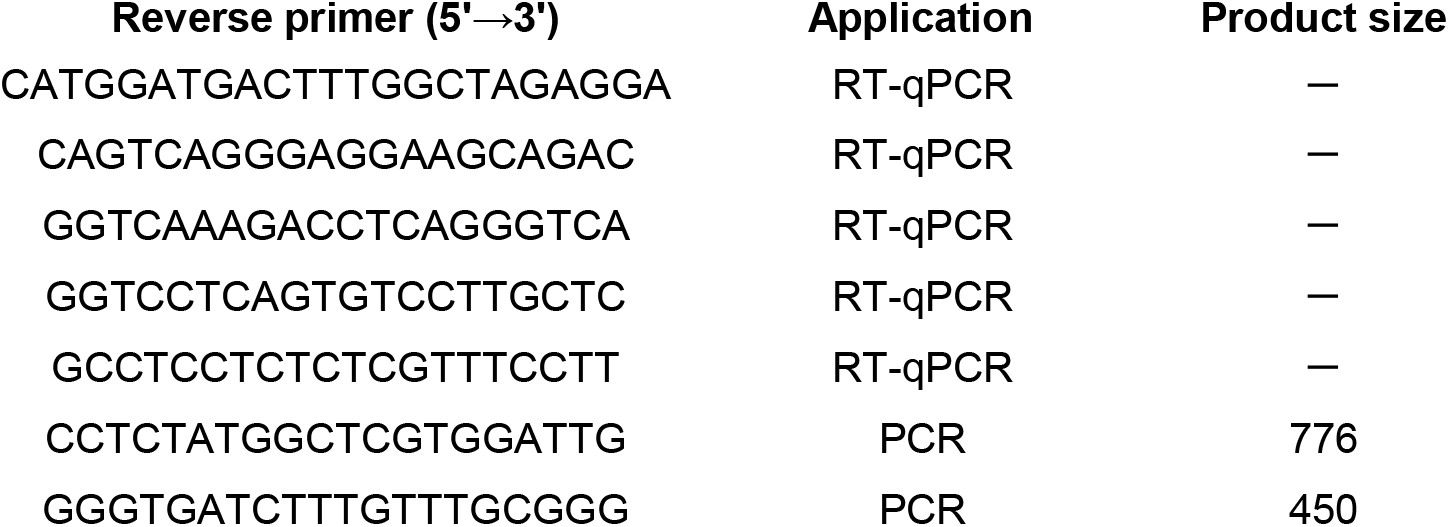
Primer sequences used for PCR and RT-qPCR analyses.

### RNA sequencing and transcriptomic analysis

RNA sequencing was performed to characterize transcriptomic alterations associated with 2-AAF exposure, multistep bladder tumor progression, and siRNA-mediated gene silencing. For all RNA sequencing analyses, total RNA was extracted using the FavorPrep Tissue Total RNA Purification Mini Kit (Favorgen Biotech Corp.). For analyses of 2-AAF-treated CNBO and siRNA-treated organoids, RNA sequencing libraries were prepared and sequenced on an Illumina platform to generate paired-end reads. Raw sequencing reads were quality filtered and aligned to the canine reference genome using HISAT2. Gene-level read counts were quantified using HTSeq, and differential expression analysis was performed using DESeq2. Differentially expressed genes (DEGs) were defined as those with an adjusted *P* value (padj) < 0.05 and an absolute log2 fold change ≥ 1. Principal component analysis (PCA) and gene set enrichment analysis (GSEA) were performed using normalized expression data. For GSEA, canine genes were converted to human orthologs and analyzed using the MSigDB Hallmark gene sets. For analyses of gene expression changes associated with bladder tumor progression, RNA sequencing libraries were prepared and sequenced on an Illumina platform to generate paired-end reads. Raw sequencing reads were mapped to the canine reference genome (CanFam6) using STAR. Gene-level read counts were quantified using RSEM. Genes with low expression (average count < 10 across all samples) were excluded from further analysis. Read counts were normalized using the trimmed mean of M-values (TMM) method. Differential expression analysis was performed using a generalized linear model likelihood ratio test (GLM-LRT) in the R package edgeR. Differentially expressed genes (DEGs) were defined based on a false discovery rate (*q*-value) < 0.05, absolute log2 fold change > 1, at least one group with counts > 50, and a coefficient of variation < 1 within each group. PCA, heatmap visualization, and GSEA were performed using normalized expression data. Canine genes were converted to human orthologs prior to GSEA.

### Immunofluorescence staining

Organoids were fixed with 4% paraformaldehyde at RT for 1 h and then immersed in 30% sucrose at 4 °C overnight. Samples were embedded in OCT compound (Sakura Finetek Japan, Tokyo, Japan) and frozen at −80 °C. Frozen blocks were sectioned into 8 µm slices using a cryostat (Leica, Wetzlar, Germany). Sections were washed three times with PBS. For detection of 2-AAF-DNA adducts, DNA was denatured by treatment with 2 M HCl at RT for 30 min before blocking. Sections were then blocked with 1.5% normal goat serum for 30 min at RT. Slides were incubated with primary antibodies listed in Table 1 at 4 °C overnight. After washing with PBS, sections were incubated with fluorescently labeled secondary antibodies and DAPI for 1 h at RT. Images were acquired using a confocal laser scanning microscope (LSM 710 NLO, Carl Zeiss, Germany). Four independent sections were analyzed for each sample. Marker-positive cells were quantified and normalized to the total number of DAPI-positive cells.

### Mouse xenograft assay

Xenograft experiments were performed with minor modifications of a previously described protocol ^69^. Male C.B-17/IcrHsd-Prkdc scid mice (5 weeks old; Japan SLC, Hamamatsu, Japan) were housed under specific pathogen-free conditions. For xenograft assays, organoid-derived cells were resuspended in a 1:1 mixture of Matrigel and organoid growth medium to a final volume of 100 μL and subcutaneously implanted into the dorsal region of mice under isoflurane anesthesia. For the primary xenograft using 2-AAF-treated CNBO-derived cells, a total of 1 × 10⁶ cells were co-injected with 5 × 10⁵ NIH 3T3 cells (ATCC, Manassas, VA, USA) per injection site. Xenograft assays using organoids established from benign xenograft tumors were performed under the same conditions. For xenograft assays using siRNA-treated organoids derived from spontaneous canine bladder carcinomas, cells were transfected with siRNA prior to implantation and injected without NIH 3T3 cells. Tumor formation was monitored, and mice were euthanized under isoflurane anesthesia at the indicated time points (21 days for xenografts derived from 2-AAF-treated CNBO and organoids established from benign xenograft tumors, and 28 days for xenografts derived from spontaneous canine bladder carcinoma organoids). Tumors were excised and processed for subsequent analyses.

### Establishment of xenograft-derived organoids

Tumors obtained from xenograft experiments were processed as previously described with minor modifications^69^. Briefly, tumor tissues were minced into small fragments and digested in 0.125 mg/mL Liberase TH (Roche Diagnostics, Indianapolis, IN, USA) at 37 °C for 30 min with gentle agitation. After centrifugation, the supernatant was removed, and the cell pellet was further dissociated using TrypLE Express at 37 °C for 5 min. The enzymatic reaction was neutralized with FBS, followed by three washes with PBS. The resulting cells were embedded in Matrigel and cultured in organoid growth medium. Organoids established from tumors with histologically benign features were defined as CBBO, whereas those derived from tumors with malignant features were defined as canine bladder carcinoma organoids (CBCO).

### Species verification of organoids

To confirm the species origin of the established organoids and assess potential contamination by mouse cells derived from xenograft hosts, species-specific PCR targeting canine and murine mitochondrial DNA was performed. Genomic DNA was extracted from organoids using the QIAamp DNA Mini Kit (QIAGEN GmbH) according to the manufacturer’s instructions. Primer sequences are listed in Table 2. A canine urothelial carcinoma cell line (AZACU) was purchased from Cosmo Bio Co., Ltd. (Tokyo, Japan) and the murine myoblast cell line C2C12 obtained from RIKEN BioResource Research Center (RIKEN BRC, Tsukuba, Ibaraki, Japan) were included as positive controls for the canine- and mouse-specific primer sets, respectively. PCR products were analyzed by agarose gel electrophoresis.

### Transmission electron microscopy

Organoids were processed for transmission electron microscopy as previously described ^69^. Organoids were initially fixed with 2.5% glutaraldehyde in 0.1 M phosphate buffer at 4 °C and subsequently post-fixed in 1% osmium tetroxide. Following dehydration through a graded ethanol series, samples were embedded in epoxy resin. Ultrathin sections (∼70 nm) were prepared and stained with uranyl acetate and lead citrate prior to observation using a transmission electron microscope.

### Proliferation and drug sensitivity assays

For the 2-AAF dose–response assay, cell viability was measured on day 6 after 2-AAF treatment using the PrestoBlue assay. In a separate proliferation assay comparing CNBO, CBBO, and CBCO, culture medium was replaced on day 4 after seeding, and cell viability was measured on day 5 using the PrestoBlue assay. For drug sensitivity assays, cells were cultured for 24 h after seeding and then treated with vinblastine (0.1, 1, and 10 nM), mitoxantrone (1, 10, and 100 ng/mL), carboplatin (1, 10, and 100 μg/mL), or lapatinib (2, 4, and 8 μM) based on a previous report^65,69^. Vehicle control cells were treated with DMSO. After 72 h of treatment, cell viability was measured as described above, normalized to vehicle-treated controls, and expressed as a percentage.

### Whole-exome sequencing and somatic variant analysis

Whole-exome sequencing (WES) data from CBBO and CBCO were analyzed using CNBO as the matched normal control. Sequence quality was assessed using FastQC, and adapter sequences and low-quality bases were trimmed using Trimmomatic. Filtered reads were aligned to the canine reference genome CanFam3.1 using BWA, and duplicate reads were removed using GATK MarkDuplicates. Somatic variants were identified using GATK Mutect2 within the exome target regions, followed by filtering with FilterMutectCalls. Detected variants were annotated using snpEff with the CanFam3.1.99 database to predict their effects on genes and transcripts. Tumor mutation burden (TMB) was calculated using pyTMB based on nonsynonymous somatic single-nucleotide variants per megabase. TMB was calculated for both all detected variants and high-quality variants that passed FilterMutectCalls. The software used in this analysis included FastQC v0.12.1, Trimmomatic v0.39, BWA v0.7.17, SAMtools v1.19, BCFtools v1.19, GATK v4.2.5.0, snpEff v5.2, and pyTMB v1.3.0. Additional downstream analyses were performed as follows. Cancer-related genes were identified with reference to the COSMIC Cancer Gene Census database. Variants were visually inspected using Integrative Genomics Viewer (IGV). Mutated genes identified by whole-exome sequencing were compared with the Cancer Gene Census (CGC) Tier 1 gene set. Genes matching the CGC database were further assessed for annotation as bladder cancer–related genes and visualized using an OncoPrint format.

### Western blot analysis

Protein lysates were prepared using a cell lysis reagent containing a protease inhibitor cocktail. Protein expression was analyzed using a Jess system (ProteinSimple, San Jose, CA, USA) according to the manufacturer’s instructions. Briefly, protein samples were mixed with fluorescent master mix and loaded into capillaries for size-based separation and immunodetection. Target proteins were incubated with primary antibodies listed in Table 1, followed by incubation with horseradish peroxidase-conjugated secondary antibodies. Chemiluminescent signals were detected and quantified using Compass for SW software (ProteinSimple). Protein expression levels were evaluated based on peak area. For MMP1, expression levels were calculated as the sum of three detected peaks.

### siRNA transfection

Cells derived from organoids were seeded at a density of 1.0 × 10⁵ cells per well in 6-well plates and cultured for 24 h. Cells were then transfected with siRNAs targeting *COL7A1* or *MMP1*, or with a negative control siRNA (Table 3), using Lipofectamine RNAiMAX (Thermo Fisher Scientific) in Opti-MEM (Thermo Fisher Scientific) at a final concentration of 40 nM. Transfection efficiency was primarily evaluated by Western blot analysis. Cells at 72 h after transfection were subjected to subsequent functional assays.

**Table 3.**
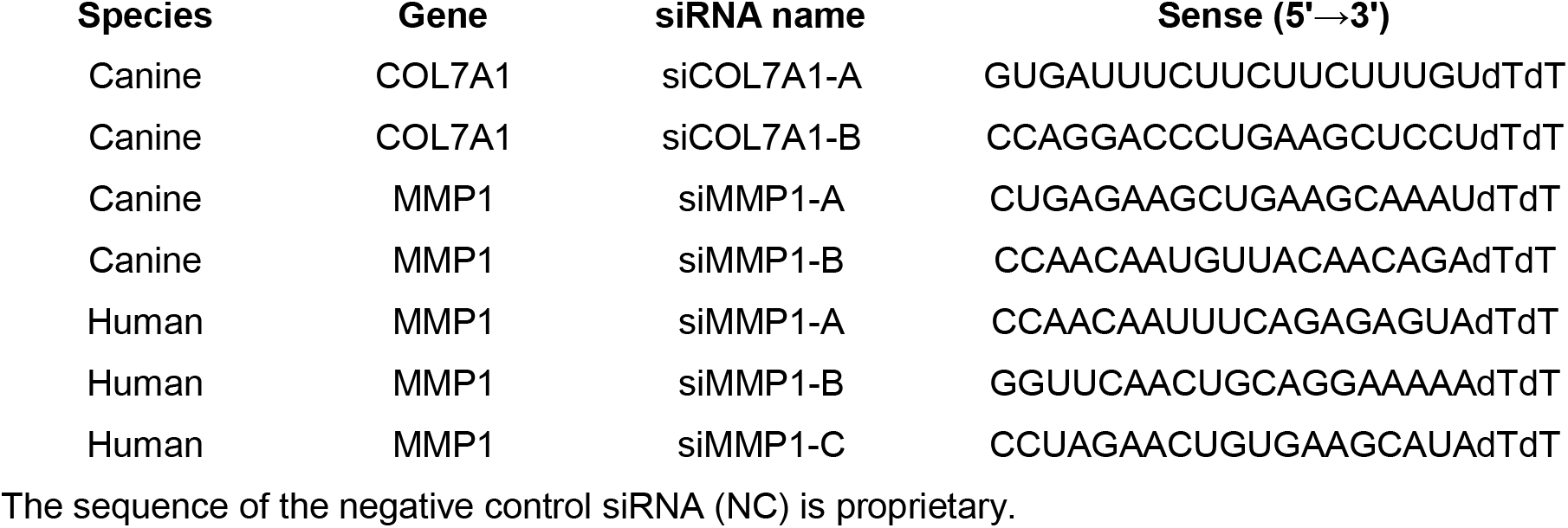
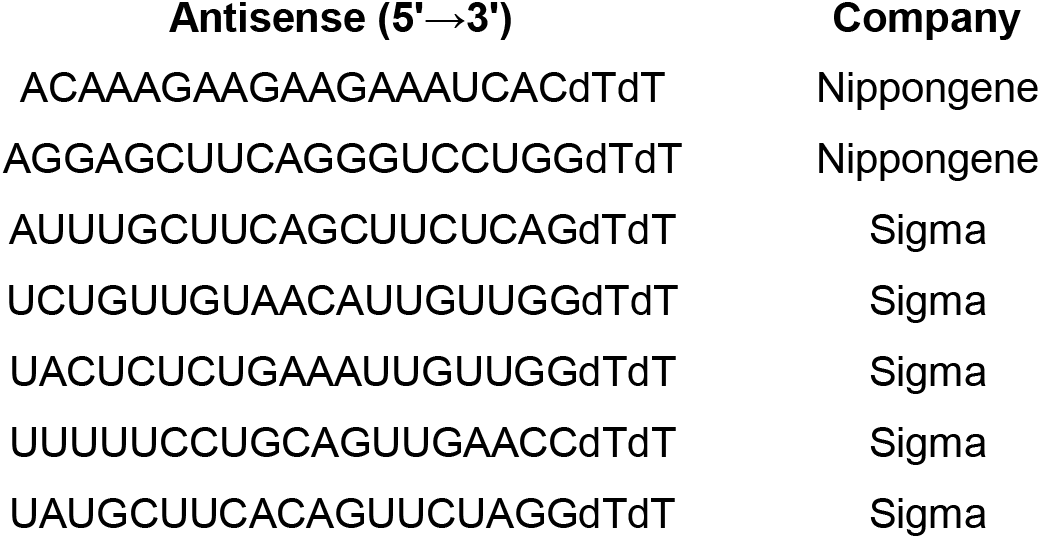
siRNA sequences used in this study.

### Cell invasion assay

Cell invasion was evaluated using a Matrigel-coated Transwell system. The membranes were coated with 2% Matrigel, and cells were seeded in the upper chamber. Medium was added to the lower chamber to induce invasion. After 24 h, non-invading cells were removed, and invaded cells were fixed with methanol for 15 min and stained with Giemsa solution (FUJIFILM Wako). The number of invaded cells was counted in six fields of view, and the average value was used for analysis.

### Human bladder cancer cell culture

Human bladder cancer cell lines 5637(RCB3676) and T24(RCB2536) were obtained from RIKEN BioResource Research Center (Ibaraki, Japan). 5637 cells were cultured in RPMI 1640 medium supplemented with 10% FBS and 1% penicillin–streptomycin. T24 cells were cultured in Dulbecco’s modified Eagle’s medium (low glucose) supplemented with 10% FBS and 1% penicillin–streptomycin. Cells were maintained under standard conditions (37 °C, 5% CO₂). The authenticity of the cell lines was confirmed by short tandem repeat (STR) profiling.

### Establishment of spontaneous canine bladder carcinoma organoids (sCBCO)

Tumor tissues were obtained from canine bladder cancer patients undergoing surgical treatment and transported at 4 °C in sterile saline. Patient information is provided in Table 4. Tumor tissues were processed for organoid culture as described above, including enzymatic digestion and Matrigel embedding. Established organoids were designated as spontaneous canine bladder carcinoma organoids (sCBCO) and used for subsequent experiments.

**Table 4.** Clinical information of dogs with spontaneous bladder cancer used for organoid establishment.

| **Sample ID** | **Age** | **Breed** | **Gender** | **Diagnosis** | **Animal hospital name** |
| --- | --- | --- | --- | --- | --- |
| sCBCO-1 | 7 y 6 m | Border Collie | Spayed female | Urothelial carcinoma | Ai animal medical center Takahagi |
| sCBCO-2 | 14 y | Maltese | Spayed female | Urothelial carcinoma | Ai sogo animal hospital |

### Ethics statement

All animal experiments, including xenograft assays using CNBO, CBBO, and sCBCO, were approved by the Institutional Animal Care and Use Committee of Tokyo University of Agriculture and Technology (approval no. R04-120) and were conducted in accordance with the ARRIVE guidelines. Spontaneous canine bladder carcinoma organoids (sCBCO) were established from tumor tissues obtained from client-owned dogs with spontaneous bladder cancer treated at veterinary hospitals. The collection and use of canine clinical samples were conducted in accordance with the guidelines of the Institutional Animal Care and Use Committee of Tokyo University of Agriculture and Technology (approval no. 0020007; approval date: 6 May 2021). Written informed consent for the use of clinical samples for research purposes was obtained from the owners of all dogs.

### Statistical analysis

All statistical analyses were performed using SigmaPlot (version 16.0, Systat Software Inc., San Jose, CA, USA) unless otherwise stated. Data are presented as mean ± standard deviation (SD). Comparisons between two groups were performed using an unpaired two-tailed Welch’s *t*-test. Comparisons among three or more groups were performed using one-way analysis of variance (ANOVA) followed by appropriate post hoc multiple-comparison tests. Non-parametric data were analyzed using the Mann–Whitney rank sum test. Sample sizes and replicate numbers for each experiment are provided in the corresponding figure legends. A *P* value < 0.05 was considered statistically significant.

## Results

### Establishment of a stepwise bladder tumor organoid model and characterization of 2-AAF–treated CNBO

Following 2-AAF treatment, CNBO were transplanted into immunodeficient mice and gave rise to benign bladder tumors. Organoids established from these tumors were designated canine benign bladder organoids (CBBO). CBBO were subsequently retransplanted into immunodeficient mice, resulting in the formation of malignant bladder carcinomas. Organoids established from these malignant tumors were designated canine bladder carcinoma organoids (CBCO). CNBO, CBBO, and CBCO were then subjected to histological, functional, transcriptomic, and genomic analyses to characterize molecular changes associated with stepwise bladder tumor progression (Fig. 1A). CNBO exhibited a cystic epithelial morphology on H&E staining and expressed the urothelial markers CK5 and UPK3A (Fig. 1B), confirming their urothelial identity. CNBO were exposed to 2-AAF at concentrations ranging from 0.01 to 10 μM for up to 6 days. Treatment with 10 μM 2-AAF resulted in marked cytotoxicity, characterized by a substantial reduction in organoid number and size, whereas no obvious cytotoxic effects were observed at concentrations of 1 μM or lower (Fig. 1C). Cell viability analysis demonstrated a dose-dependent decrease in viability following 2-AAF exposure, with an estimated IC50 value of 8.4 μM after 6 days of treatment (Fig. 1D). Based on these findings, 1 μM 2-AAF was selected for subsequent carcinogen exposure experiments, as it induced cellular responses without causing substantial cytotoxicity. To investigate the molecular response of CNBO to 2-AAF exposure, we analyzed the expression of representative metabolic and stress-response genes. After 24 h of 2-AAF treatment, expression levels of *CYP4B1*, *HMOX1*, and *NQO1* were significantly increased (Fig. 1E). RNA-seq analysis identified enrichment of pathways related to glutathione metabolism and detoxification in 2-AAF-treated CNBO (Fig. 1F). Consistent with this, multiple glutathione metabolism–associated genes, including *GPX2*, *GPX3*, *GSS*, *GCLC*, *GGT1*, *GSR*, and *MGST1*, were upregulated following 2-AAF exposure (Fig. 1G). After 3 days of 2-AAF exposure, immunofluorescence analysis revealed significantly increased AAF-DNA adduct formation and γH2AX positivity in CNBO compared with vehicle-treated controls, indicating enhanced DNA damage (Fig. 1H, I). These findings suggest that 2-AAF induces early molecular changes associated with bladder carcinogenesis in CNBO.

**Figure 1.**
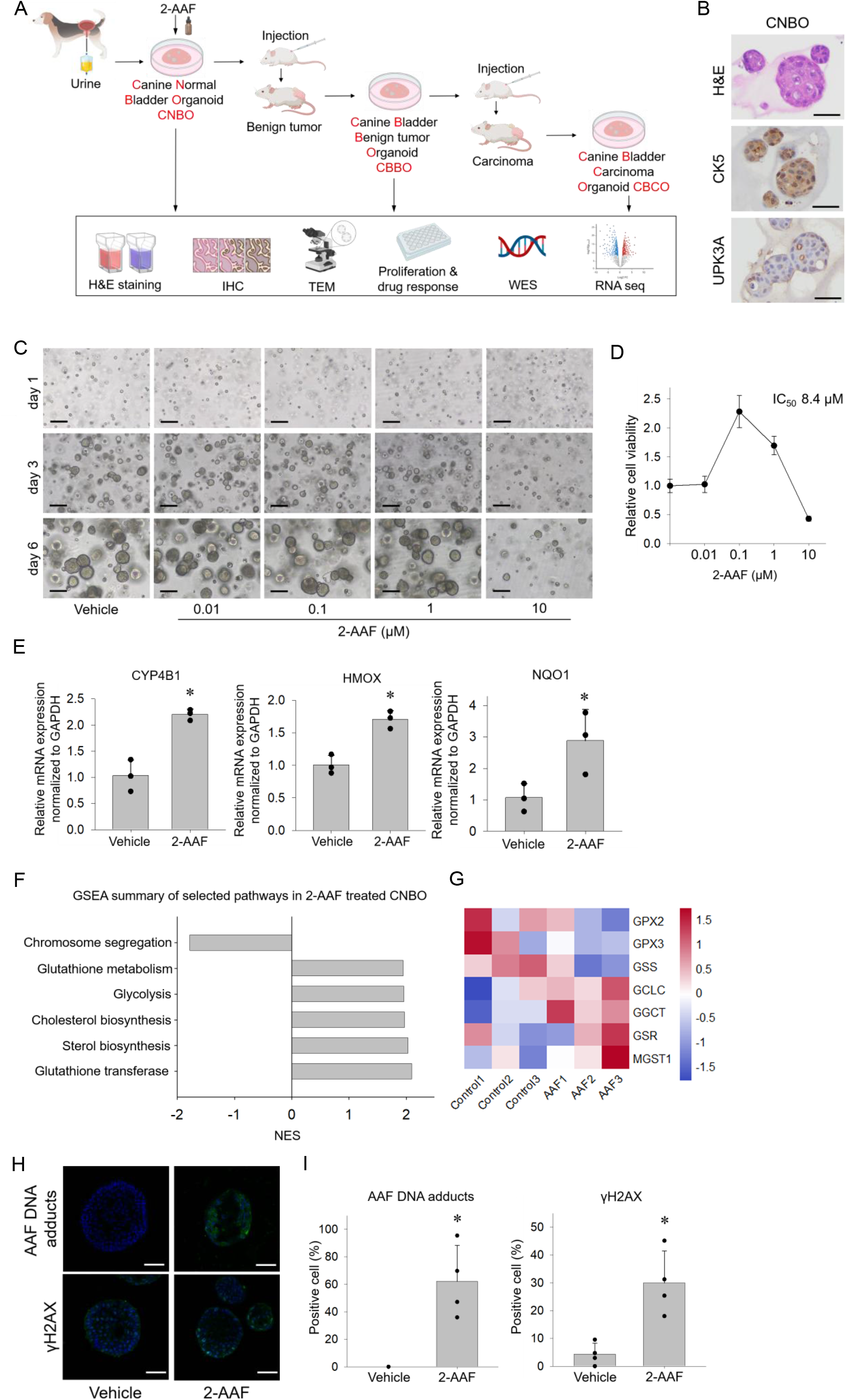
Establishment of a stepwise bladder tumor organoid model and characterization of 2-AAF–treated normal canine bladder organoids (CNBO). (A) Schematic diagram of the stepwise bladder tumor organoid model. CNBO were treated with 1 μM 2-AAF for 6 days and transplanted into immunodeficient mice. Twenty-one days after transplantation, the resulting benign bladder tumors were collected, and organoids established from these lesions were designated canine benign bladder organoids (CBBO). CBBO were subsequently retransplanted into immunodeficient mice, and the resulting bladder carcinomas were harvested 21 days later. Organoids established from these malignant lesions were designated canine bladder carcinoma organoids (CBCO). Organoids at each stage (CNBO, CBBO, and CBCO) underwent histological, immunohistochemical, ultrastructural, functional, transcriptomic, and genomic analyses. (B) Representative Hematoxylin and Eosin (H&E) and immunohistochemical staining images of CNBO for CK5 and UPK3A. Scale bar, 40 μm. (C) Representative bright-field images of CNBO treated with the indicated concentrations of 2-AAF for 1, 3, and 6 days. Scale bar, 300 μm. (D) Cell viability of CNBO following treatment with the indicated concentrations of 2-AAF was evaluated by Presto Blue assay. Data are presented as mean ± SD (n = 4). (E) The effects of 2-AAF (1 µM, 24 h) on the expression of xenobiotic metabolism- and oxidative stress-related genes, *CYP4B1*, *HMOX1*, and *NQO1* in CNBO were evaluated by quantitative PCR and expressed as mean ± SD. n=4. \**P*<0.05 vs. Control. (F) Gene Set Enrichment Analysis (GSEA) using RNA-sequencing data in CNBO after 24 h of 1 µM 2-AAF treatment. (G) Heatmap of glutathione metabolism–associated gene expression in vehicle- and 2-AAF-treated CNBO. (H) Representative immunofluorescence images of AAF-DNA adducts and γH2AX expression in CNBO after 72 h of 2-AAF treatment. Scale bar, 50 μm. (I) Quantification of AAF-DNA adduct–positive and γH2AX-positive cells in CNBO after 72 h of 2-AAF treatment. Data are presented as mean ± SD from four independent sections. \**P* < 0.05 vs. Vehicle.

### Establishment and characterization of canine benign bladder organoids (CBBO) and canine bladder cancer organoids (CBCO)

To evaluate the tumorigenic potential of 2-AAF-treated CNBO, organoids were transplanted into immunodeficient mice. Tumors formed and were harvested on day 21 (Fig. 2A). The isolated tumor tissues exhibited a cystic architecture lined by proliferating urothelial cells, with focal papillary epithelial proliferation. Nuclear atypia was mild, and no invasive growth was observed. Based on these findings, the tumor tissues were diagnosed as benign urothelial tumors (Fig. 2B). We next established canine benign bladder organoids (CBBO) from the benign tumor tissues and confirmed their continuous propagation in vitro (Fig. 2C). CBBO were subsequently retransplanted into immunodeficient mice, and tumors were successfully generated and harvested 21 days after transplantation for further histopathological analysis (Fig. 2D). The tumor tissues showed atypical epithelial proliferation with nuclear atypia, architectural disorganization, and loss of cell cohesion. Invasive growth into the surrounding stroma was also observed. Based on these histopathological features, they were diagnosed as carcinoma (Fig. 2E). We next established organoids from the malignant tumors, designated them as canine bladder carcinoma organoids (CBCO), and used them for subsequent analyses (Fig. 2F). Histological and immunohistochemical analyses demonstrated that both CBBO and CBCO retained epithelial morphology and continued to express the urothelial markers CK5 and UPK3A (Fig. 2G). Transmission electron microscopy showed that CNBO retained ultrastructural features of normal urothelium, including uniform cellular morphology and a well-defined intercellular lateral boundary (ILB). In contrast, CBCO exhibited nuclear enlargement, formation of multiple lumen-like structures, and an incomplete and discontinuous ILB accompanied by cytoplasmic degenerative changes (Fig. 2H). Analysis of organoid growth revealed that the proliferative capacity of CBBO and CBCO was significantly reduced compared with that of CNBO, while no significant difference was observed between CBBO and CBCO (Fig. 2I, J). To evaluate changes in drug responsiveness during tumor progression, the sensitivities of CBBO and CBCO to anticancer drugs were compared with those of CNBO. Both CBBO and CBCO exhibited increased resistance to vinblastine, lapatinib, and mitoxantrone compared with CNBO, whereas sensitivity to carboplatin remained largely unchanged among the organoid lines (Fig. 2K).

**Figure 2.**
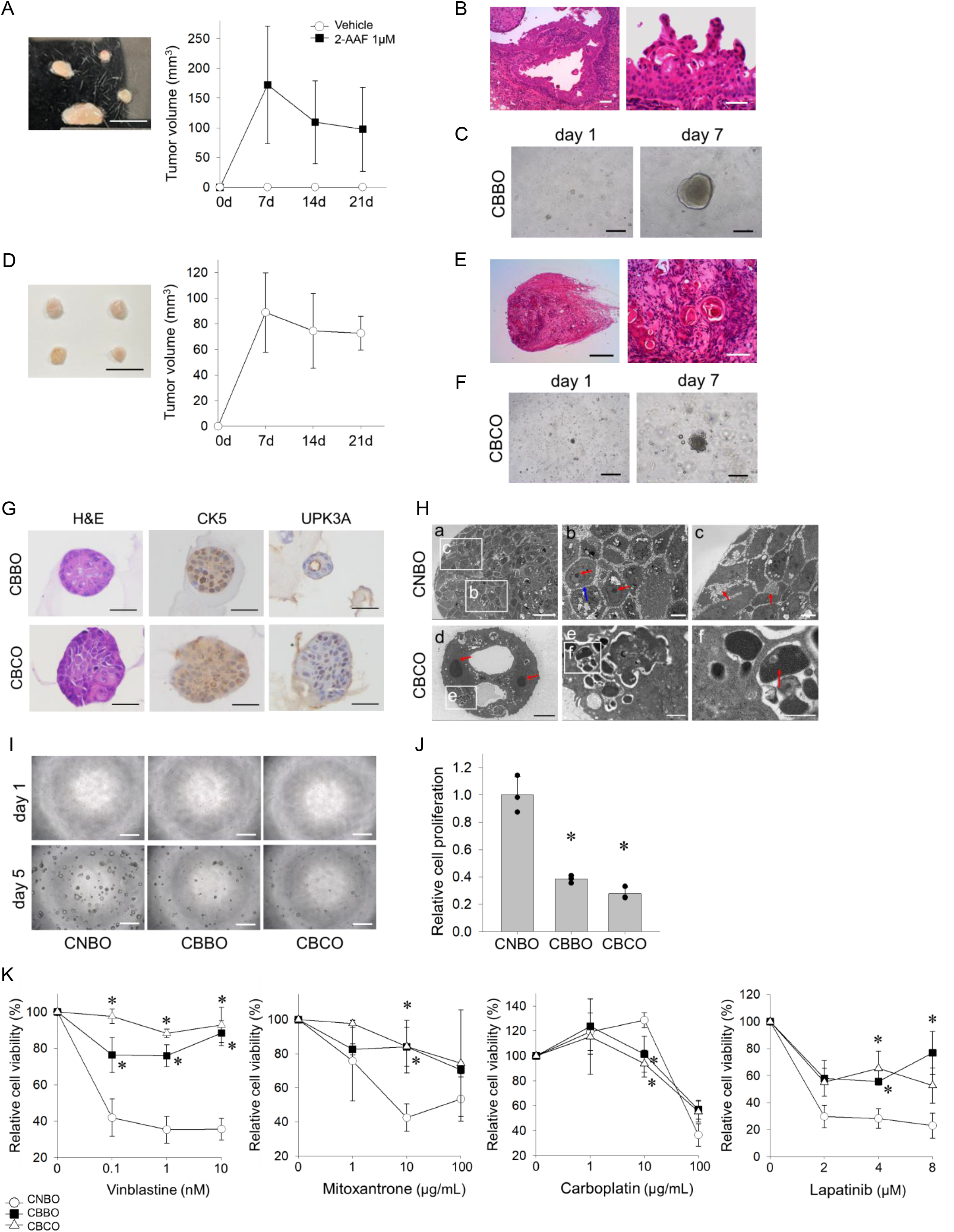
Generation and characterization of CBBO and CBCO. (A) Gross images of mass lesions formed following transplantation of 2-AAF-treated CNBO into immunodeficient mice (left). Tumor volume was measured longitudinally after transplantation (right). Data are presented as mean ± SD (n = 4). Scale bar, 10 mm. (B) H&E-stained images of lesions formed after transplantation of 2-AAF-treated CNBO into immunodeficient mice. Scale bars, 100 µm (left) and 50 µm (right). (C) Bright-field images on Day 1 and Day 7 after CBBO generation. Scale bar, 100 µm. (D) Gross images of mass lesions formed following transplantation of CBBO into immunodeficient mice (left). Tumor volume was measured longitudinally after transplantation (right). Data are presented as mean ± SD (n = 4). Scale bar, 10 mm. (E) H&E-stained images of lesions formed after transplantation of CBBO into immunodeficient mice. Scale bars, 50 µm (left) and 100 µm (right). (F) Bright-field images on Day 1 and Day 7 after CBCO generation. Scale bar, 100 µm. (G) Representative H&E and immunohistochemical staining images of CBBO and CBCO for CK5 and UPK3A. Scale bar, 40 µm. (H) Transmission electron microscopy (TEM) images of CNBO and CBCO. (a) Low-magnification image of CNBO. (b) Higher-magnification image of CNBO showing nuclei and nucleoli. (c) Higher-magnification image of CNBO showing the intercellular lateral boundary (ILB). (d) Low-magnification image of CBCO showing multiple lumen-like structures. (e) Higher-magnification image of CBCO showing the ILB. (f) Higher-magnification image of CBCO showing the ILB and cytoplasmic changes. Scale bars, 20 µm (a), 5 µm (b, c), 80 µm (d), 1 µm (e), and 500 nm (f). (I) Representative bright-field images of CNBO, CBBO, and CBCO from the cell proliferation assay at Days 1 and 5. For each condition, 500 cells were suspended in Matrigel, seeded into 96-well plates, and photographed at the indicated time points. Scale bar, 50 µm. (J) Quantification of the cell proliferation assay using PrestoBlue assay at Day 5. Data are presented as mean ± SD (n = 3). (K) Drug sensitivity assays in CNBO, CBBO, and CBCO at 72 h after treatment with vinblastine (0.1-10 nM), mitoxantrone (1-100 ng/ml), carboplatin (1-100 µg/ml), or lapatinib (2-8 µM). Data are presented as mean ± SD (n = 3). \**P* < 0.05 vs. CNBO.

### Genomic characterization of CBBO and CBCO by whole-exome sequencing

Whole-exome sequencing using CNBO as the reference identified 131 mutations in CBBO and 69 mutations in CBCO, predominantly consisting of missense variants with fewer frameshift and splice-site mutations (Fig. 3A). Tumor mutational burden (TMB) was relatively low in both organoids, with values of 0.94 Mut/Mb in CBBO and 0.63 Mut/Mb in CBCO (Fig. 3B). Nine mutated genes were shared between CBBO and CBCO (Fig. 3C). IGV analysis showed an increase in mutant reads for *DYRK1A* and *JARID2* from CNBO to CBBO and CBCO (Fig. 3D). Although several mutations were detected in cancer-associated genes listed in the Cancer Gene Census (CGC) Tier 1 database, none were annotated as bladder cancer–related genes (Fig. 3E). Collectively, these findings indicate that tumor progression in this model occurs in the context of relatively limited genomic alterations.

**Figure 3.**
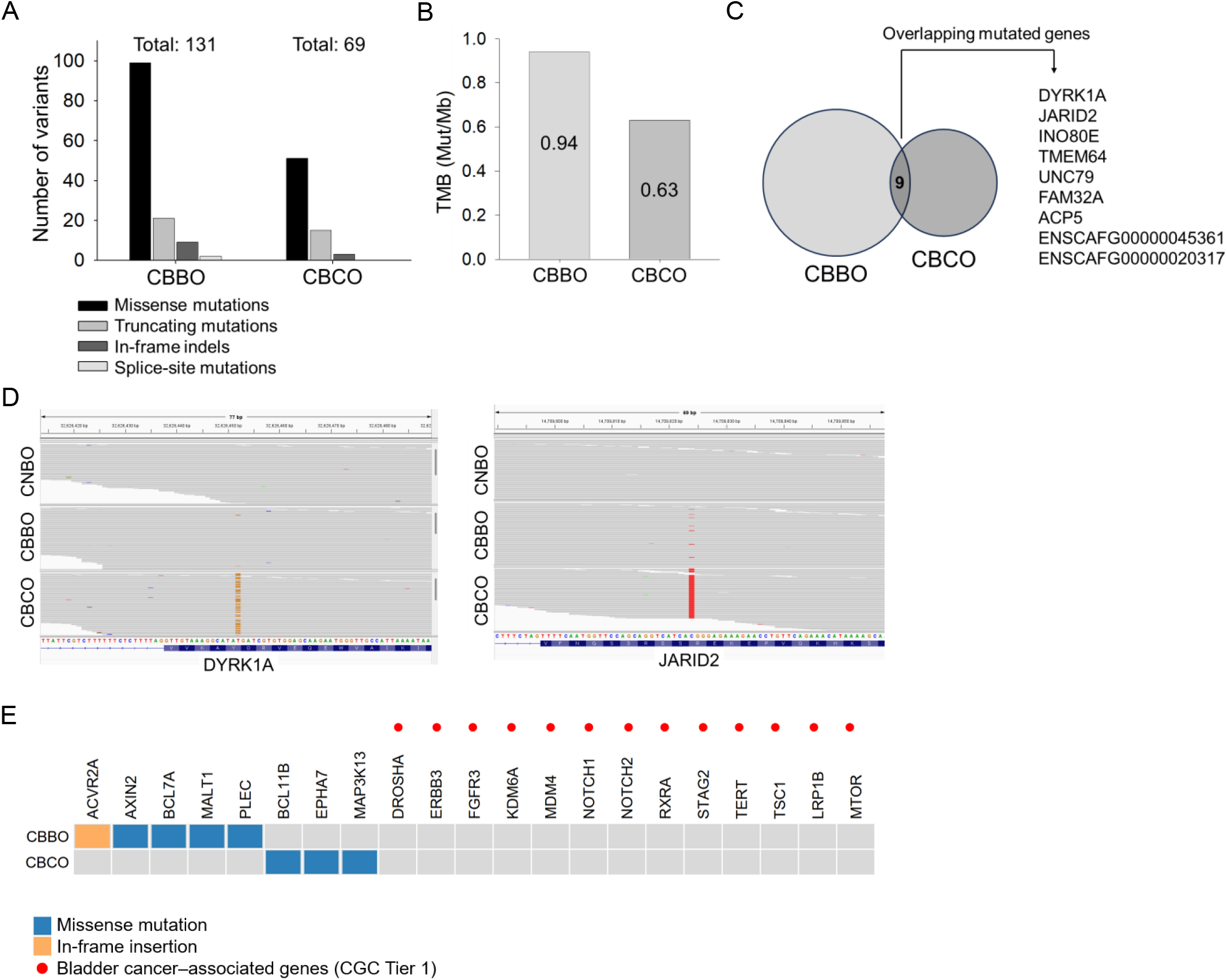
Whole-exome sequencing analysis of CBBO and CBCO. (A) Overview of somatic mutations identified in CBBO and CBCO. Variants predicted to have high or moderate impact by SnpEff are classified according to mutation type. (B) Comparison of tumor mutation burden (TMB). TMB was calculated using high-confidence nonsynonymous somatic single-nucleotide variants that passed FilterMutectCalls (PASS). (C) Overlap of somatically mutated genes in CBBO and CBCO. (D) Read alignments at mutation loci of *DYRK1A* and *JARID2* in CNBO, CBBO, and CBCO. (E) OncoPrint representation of somatic mutations in Cancer Gene Census (CGC) Tier 1 genes in CBBO and CBCO.

### Progressive activation of EMT-associated molecular programs during bladder tumor organoid progression

To investigate transcriptional changes during tumor progression, RNA sequencing was performed using CNBO, CBBO, and CBCO. Differential expression analysis identified widespread transcriptional alterations, with 263 upregulated and 430 downregulated genes in CBBO versus CNBO, 530 upregulated and 230 downregulated genes in CBCO versus CBBO, and 450 upregulated and 368 downregulated genes in CBCO versus CNBO (Fig. 4A). Gene set enrichment analysis (GSEA) revealed consistent enrichment of the epithelial–mesenchymal transition (EMT) pathway during tumor progression in both the CNBO–CBBO and CBBO–CBCO comparisons (Fig. 4B). Heatmap analysis of the top 30 differentially expressed genes identified *COL7A1* and *MMP1* as candidates of EMT-associated genes (Fig. 4C). Expression of both *COL7A1* and *MMP1* was progressively upregulated during tumor progression, with the highest levels observed in CBCO (Fig. 4D). These transcriptional changes were further evaluated at the protein level. Immunofluorescence analysis revealed a significant increase in COL7A1, MMP1, and vimentin expression and a significant decrease in E-cadherin expression during tumor progression (Fig. 4E, F). Western blot analysis demonstrated progressive upregulation of COL7A1 and vimentin during tumor progression, whereas MMP1 expression was significantly increased in both CBBO and CBCO relative to CNBO (Fig. 4G, H). Overall, these findings indicate progressive activation of EMT-related molecular programs during bladder tumor organoid progression. Based on these results, *COL7A1* and *MMP1* were selected for further functional analyses.

**Figure 4.**
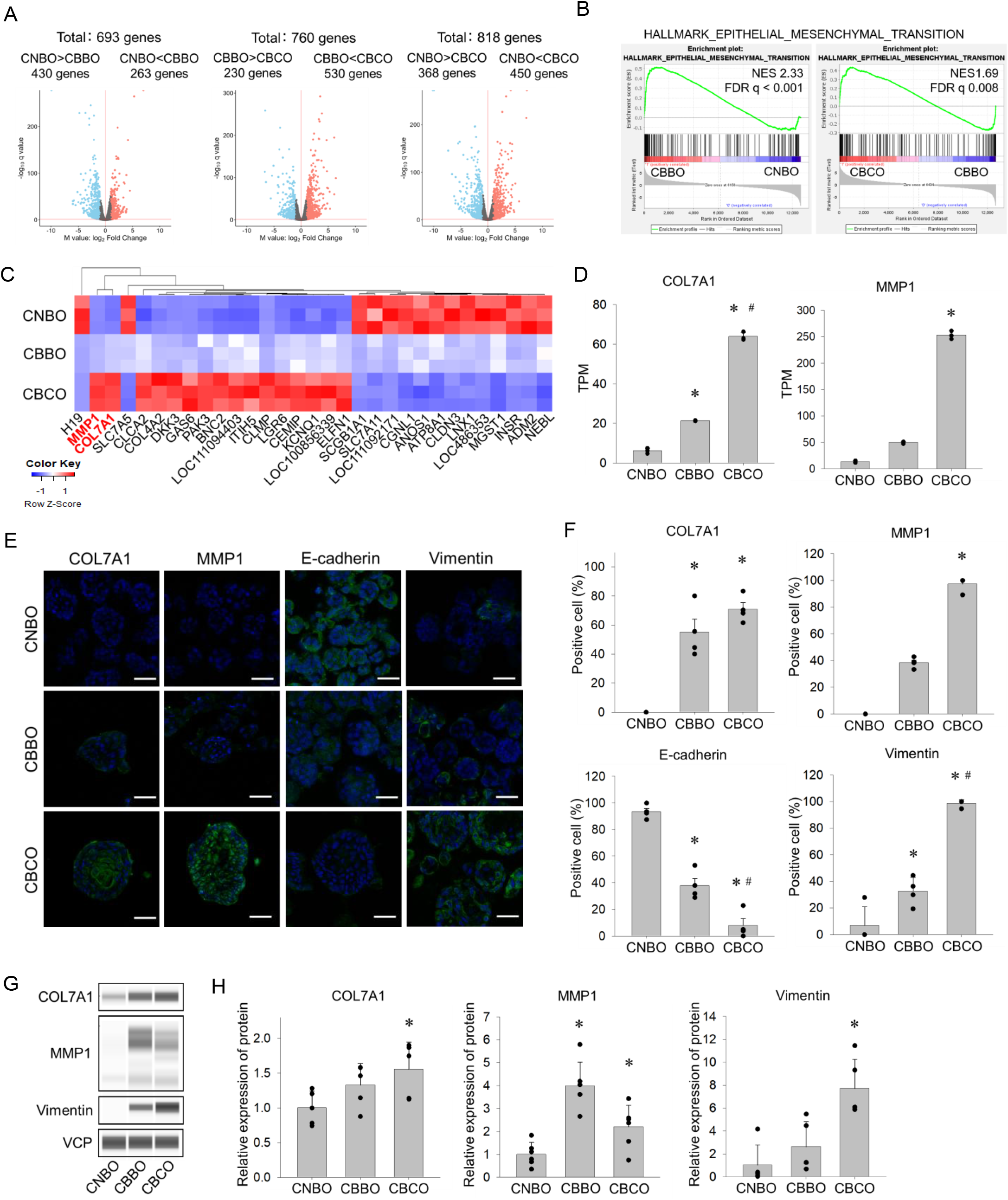
Progressive EMT activation and upregulation of *COL7A1* and *MMP1* during bladder tumor organoid progression. (A) Volcano plots of differentially expressed genes in CBBO versus CNBO, CBCO versus CBBO, and CBCO versus CNBO. (B) GSEA of the Hallmark epithelial–mesenchymal transition (EMT) gene set in CNBO–CBBO and CBBO–CBCO comparisons. (C) Heatmap of the top 30 differentially expressed genes across CNBO, CBBO, and CBCO after gene-wise Z-score normalization. *COL7A1* and *MMP1* are highlighted in red. (D) Expression analysis of *COL7A1* and *MMP1* in CNBO, CBBO, and CBCO by RNA sequencing (TPM). (E) Representative immunofluorescence images of COL7A1, MMP1, E-cadherin, and vimentin in CNBO, CBBO, and CBCO. Nuclei were counterstained with DAPI. Scale bars, 50 µm. (F) Quantification of COL7A1-, MMP1-, E-cadherin-, and vimentin-positive cells in CNBO, CBBO, and CBCO. Data are presented as mean ± SD from four independent sections. (G) Protein expression of COL7A1, MMP1, and vimentin in CNBO, CBBO, and CBCO determined by Western blot analysis. VCP was used as a loading control. (H) Quantification of COL7A1, MMP1, and vimentin protein expression in CNBO, CBBO, and CBCO. Data are presented as mean ± SD (n = 4 for COL7A1, n = 6 for MMP1, and n = 5 for vimentin). \**P* < 0.05 vs. CNBO; #*P* < 0.05 vs. CBBO.

### Functional analysis of COL7A1 and MMP1 in canine bladder cancer organoids (CBCO)

To investigate the functions of *COL7A1* and *MMP1* identified by RNA-seq analysis, knockdown experiments were performed in CBCO using two independent siRNAs targeting each gene (Fig. 5A). Efficient knockdowns of *COL7A1* and *MMP1* following siRNA transfection were confirmed by Western blot analysis, which demonstrated a significant reduction in the expression of each target protein compared with the control siRNA group (Fig. 5B, C). Next, the effects of gene knockdown on organoid growth were evaluated. Silencing of either *COL7A1* or *MMP1* significantly inhibited CBCO proliferation, confirming the growth-promoting roles of both genes (Fig. 5D, E). In addition, silencing of either *COL7A1* or *MMP1* significantly suppressed the invasive capacity of CBCO (Fig. 5F, G). Collectively, these findings indicate that *COL7A1* and *MMP1* are involved in the regulation of proliferation and invasion in CBCO.

**Figure 5.**
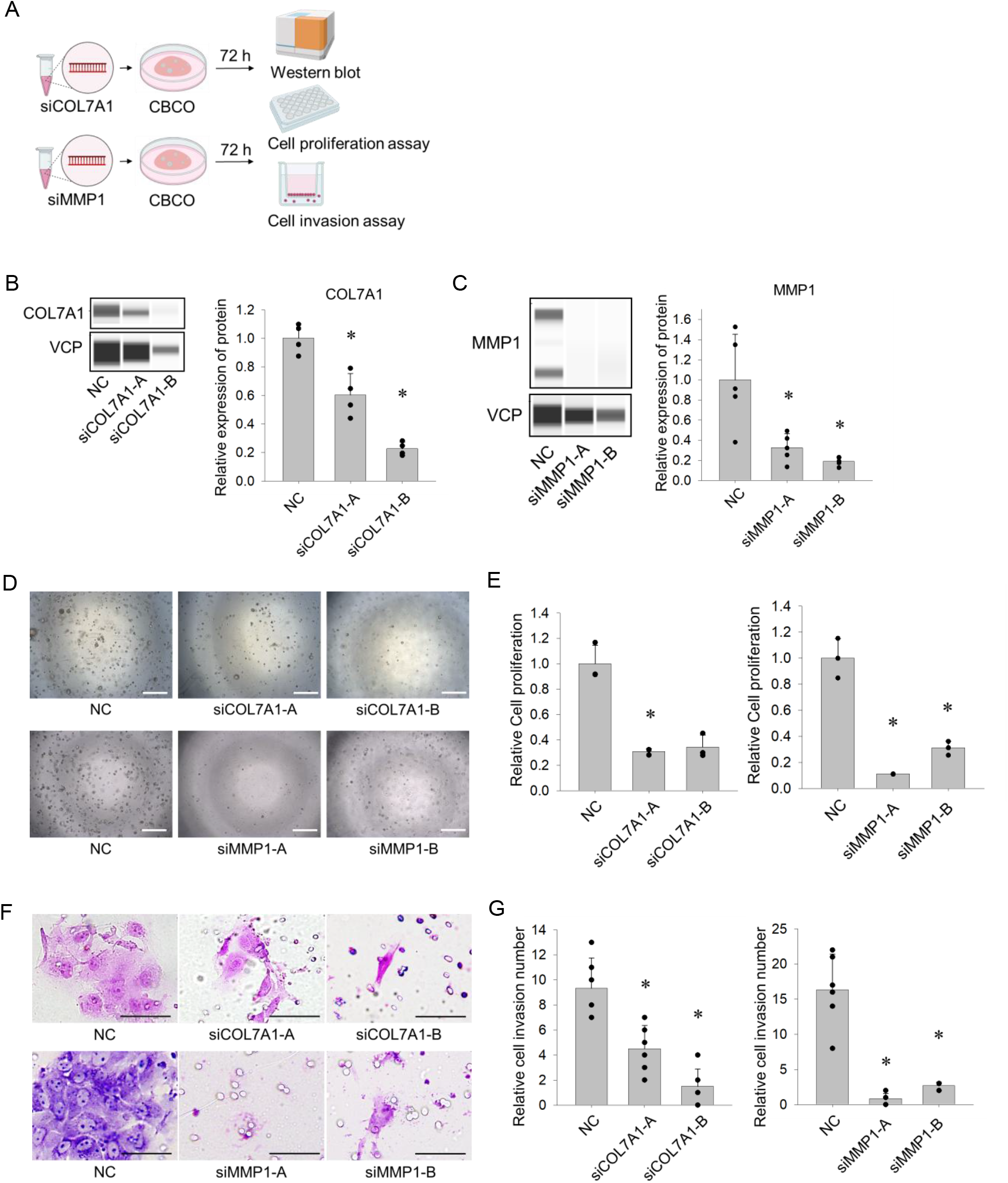
Functional analysis of COL7A1 and MMP1 knockdown in CBCO. (A) Schematic illustration of experimental design. CBCO were transfected with siRNAs targeting *COL7A1* or *MMP1* and analyzed by Western blot, cell proliferation, and cell invasion assays 72 h after transfection. (B) Western blot analysis and quantification of COL7A1 expression in CBCO transfected with NC, siCOL7A1-A, or siCOL7A1-B. VCP was used as a loading control. Data are presented as mean ± SD (n = 4). (C) Western blot analysis and quantification of MMP1 expression in CBCO transfected with NC, siMMP1-A, or siMMP1-B. VCP was used as a loading control. Data are presented as mean ± SD (n = 5). (D) Representative bright-field images from the cell proliferation assay of CBCO transfected with NC or siRNAs targeting *COL7A1* or *MMP1* on Day 7. Scale bar, 50 µm. (E) Quantification of cell proliferation in CBCO transfected with NC or siRNAs targeting *COL7A1* or *MMP1*. Data are presented as relative values normalized to NC (n = 3). (F) Representative bright-field images from the cell invasion assay of CBCO transfected with NC or siRNAs targeting *COL7A1* or *MMP1*. Scale bar, 100 µm. (G) Quantification of invaded cell numbers in CBCO transfected with NC or siRNAs targeting *COL7A1* or *MMP1*. Data are presented as mean ± SD from six independent fields. \**P* < 0.05 vs. NC.

### Functional validation of COL7A1 and MMP1 in spontaneous canine bladder carcinoma organoids

To determine whether the functions of *COL7A1* and *MMP1* identified in chemically induced CBCO are also conserved in spontaneous canine bladder carcinoma organoids (sCBCO), knockdown experiments were performed using bladder cancer organoid lines established from two dogs with spontaneous bladder cancer (sCBCO-1 and sCBCO-2) (Fig. 6A). Western blot analysis confirmed efficient knockdown of *COL7A1* by both siCOL7A1-A and siCOL7A1-B, and of *MMP1* by both siMMP1-A and siMMP1-B in sCBCO-1 and sCBCO-2 (Fig. 6B, C). Furthermore, knockdown of either *COL7A1* or *MMP1* reduced cell proliferation in both spontaneous bladder cancer organoid lines, with significant growth inhibition observed in most conditions (Fig. 6D).

**Figure 6.**
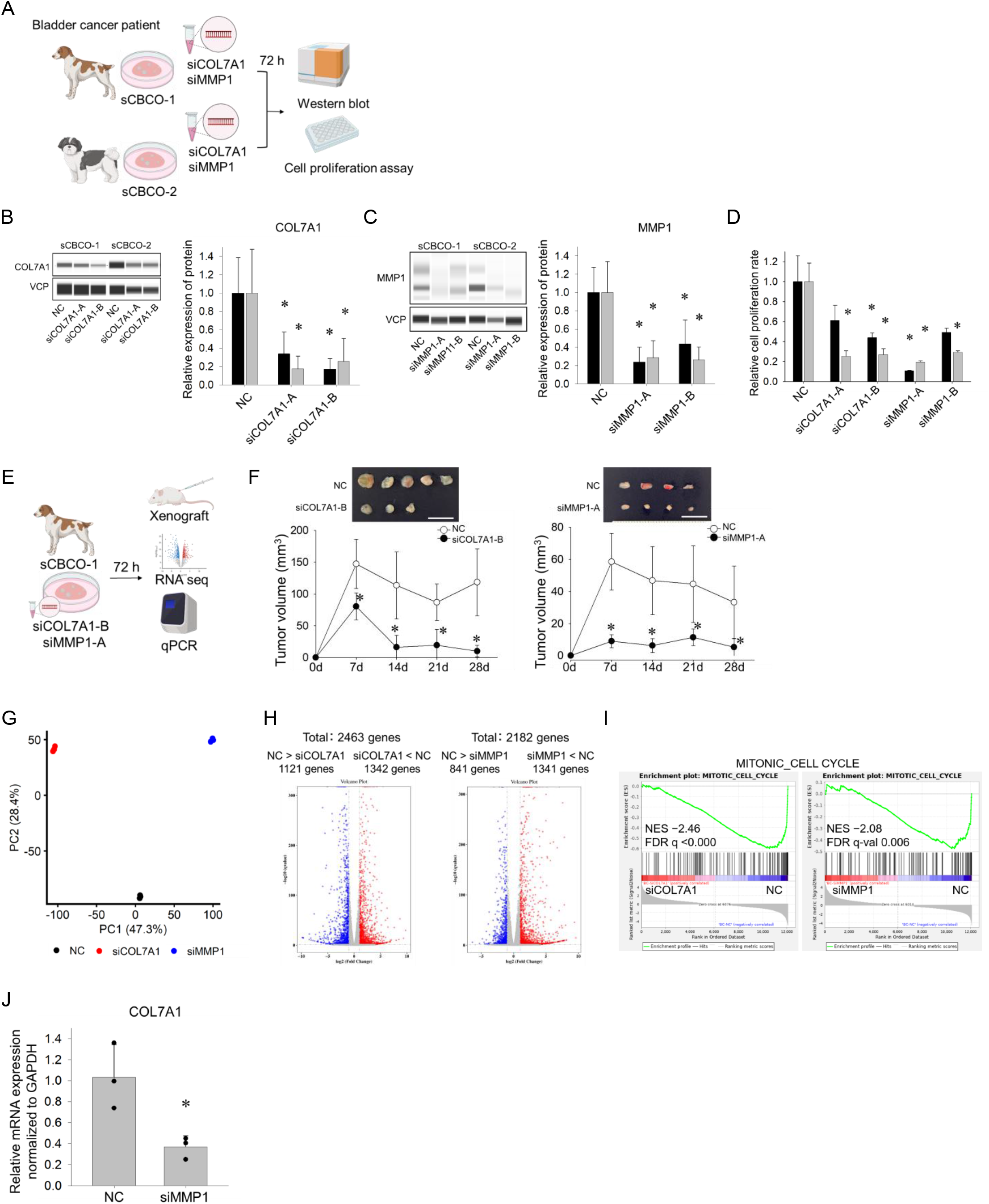
Functional validation of COL7A1 and MMP1 in spontaneous canine bladder cancer organoids. (A) Schematic illustration of experimental design. Two sCBCO lines (sCBCO-1 and sCBCO-2) were transfected with siRNAs targeting *COL7A1* or *MMP1* and analyzed by Western blot and cell proliferation assays 72 h after transfection. (B) Western blot analysis and quantification of COL7A1 expression in sCBCOs transfected with NC, siCOL7A1-A, or siCOL7A1-B. VCP was used as a loading control. Data are presented as mean ± SD (n = 5). (C) Western blot analysis and quantification of MMP1 expression in sCBCOs transfected with NC, siMMP1-A, and siMMP1-B. VCP was used as a loading control. Data are presented as mean ± SD (n = 5 for sCBCO-1 and n = 6 for sCBCO-2). (D) Quantification of cell proliferation in sCBCOs transfected with NC or siRNAs targeting *COL7A1* or *MMP1*. Data are presented as relative values normalized to NC (n = 3). (E) Schematic illustration of the experimental workflow. sCBCO-1 organoids transfected with siCOL7A1-B or siMMP1-A were transplanted into immunodeficient mice, and RNA-seq and qPCR analyses were performed in parallel. (F) Tumor growth of sCBCO-1 xenografts following transfection with siCOL7A1-B or siMMP1-A. Gross images of tumors harvested at day 28 are shown above the graphs. Scale bar, 10 mm. Tumor volumes are presented as mean ± SD (n = 5). (G) PCA of RNA-seq data from sCBCO-1 transfected with siCOL7A1-B or siMMP1-A. Each point represents an individual sample. (H) Volcano plots of RNA-seq data from sCBCO-1 transfected with siCOL7A1-B or siMMP1-A. Left: siCOL7A1-B vs. NC; right: siMMP1-A vs. NC. (I) GSEA of the Gene Ontology biological process gene set “MITOTIC CELL CYCLE” in sCBCO-1 transfected with siCOL7A1-B or siMMP1-A. (J) qPCR analysis of COL7A1 expression in sCBCO-1 transfected with siMMP1-A. Expression levels were normalized to GAPDH and are presented as relative values compared with NC (mean ± SD, n = 3). \**P* < 0.05 vs. NC.

To investigate the functional roles and molecular mechanisms of COL7A1 and MMP1, sCBCO-1 organoids were transfected with the most effective siRNAs (siCOL7A1-B and siMMP1-A). At 72 h after transfection, organoids were subjected to xenograft transplantation, RNA sequencing, and qPCR analyses (Fig. 6E). Xenograft experiments demonstrated that knockdowns of either *COL7A1* or *MMP1* significantly inhibited *in vivo* tumor growth compared with the negative control group (Fig. 6F). In RNA sequencing analysis, principal component analysis (PCA) of the RNA-seq data showed clear separation among the negative control, siCOL7A1-B, and siMMP1-A groups (Fig. 6G). Volcano plot analysis also revealed widespread transcriptional alterations following knockdown of either COL7A1 or MMP1 compared with the negative control (Fig. 6H). Gene set enrichment analysis identified multiple pathways related to cell cycle progression, mitosis, and chromosome segregation that were commonly downregulated following *COL7A1* or *MMP1* knockdown (Table 5). In particular, the mitotic cell cycle gene set showed significant negative enrichment in both siCOL7A1-B and siMMP1-A groups (Fig. 6I). Finally, qPCR analysis revealed that *COL7A1* expression was significantly reduced following *MMP1* knockdown (Fig. 6J). Collectively, these findings indicate that *COL7A1* and *MMP1* contribute to the proliferative capacity of spontaneous canine bladder cancer organoids and are associated with the regulation of mitosis-related gene expression. Furthermore, the reduction in *COL7A1* expression following *MMP1* knockdown suggests a potential regulatory relationship between *MMP1* and *COL7A1*.

**Table 5.** Commonly negatively enriched gene sets following COL7A1 or MMP1 knockdown in sCBC.

| **Gene set** | **NES (siCOL7A1-B)** | **FDR q-value** |
| --- | --- | --- |
| Mitotic cell cycle | -2.46 | <0.001 |
| G2/M transition of mitotic cell cycle | -1.99 | 0.015 |
| Mitotic cytokinesis | -2.06 | 0.006 |
| Mitotic spindle organization | -2.34 | <0.001 |
| Mitotic spindle assembly checkpoint signaling | -2.23 | <0.001 |
| Mitotic metaphase chromosome alignment | -1.94 | 0.023 |
| Chromosome segregation | -2.43 | <0.001 |
| Mitotic sister chromatid segregation | -2.27 | <0.001 |
| Chromosome centromeric region | -2.18 | 0.001 |
| Kinetochore | -2.26 | <0.001 |
| DNA replication initiation | -1.94 | 0.023 |

**O-1.**
| **NES (siMMP1-A)** | **FDR q-value** |
| --- | --- |
| -2.08 | 0.006 |
| -2.13 | 0.004 |
| -1.96 | 0.024 |
| -2.45 | <0.001 |
| -2.12 | 0.004 |
| -2.03 | 0.011 |
| -2.50 | <0.001 |
| -2.70 | <0.001 |
| -2.14 | 0.004 |
| -2.37 | <0.001 |
| -2.00 | 0.015 |

### Functional validation of MMP1 in human bladder cancer

Given the functional importance of MMP1 in regulating bladder cancer organoid growth and its potential regulatory relationship with COL7A1, we next evaluated MMP1 expression in human bladder cancer using TCGA-BLCA data. *MMP1* expression was significantly elevated in the basal/squamous subtype compared with normal bladder tissue (Fig. 7A). To further investigate the functional role of *MMP1* in human bladder cancer, siRNA-mediated knockdown of *MMP1* was performed in human bladder cancer cell lines, T24 and 5637 cells (Fig. 7B). Efficient MMP1 knockdown was confirmed by Western blot analysis, and suppression of *MMP1* significantly reduced cell proliferation in both 5637 and T24 cells (Fig. 7C–E). Collectively, these findings support the cross-species relevance of MMP1 identified using the stepwise organoid model. The proposed molecular events associated with the stepwise progression from CNBO to CBBO and CBCO, including DNA damage responses, proliferative signaling, malignant progression, and EMT activation involving *MMP1* and *COL7A1*, are summarized in Fig. 7F.

**Figure 7.**
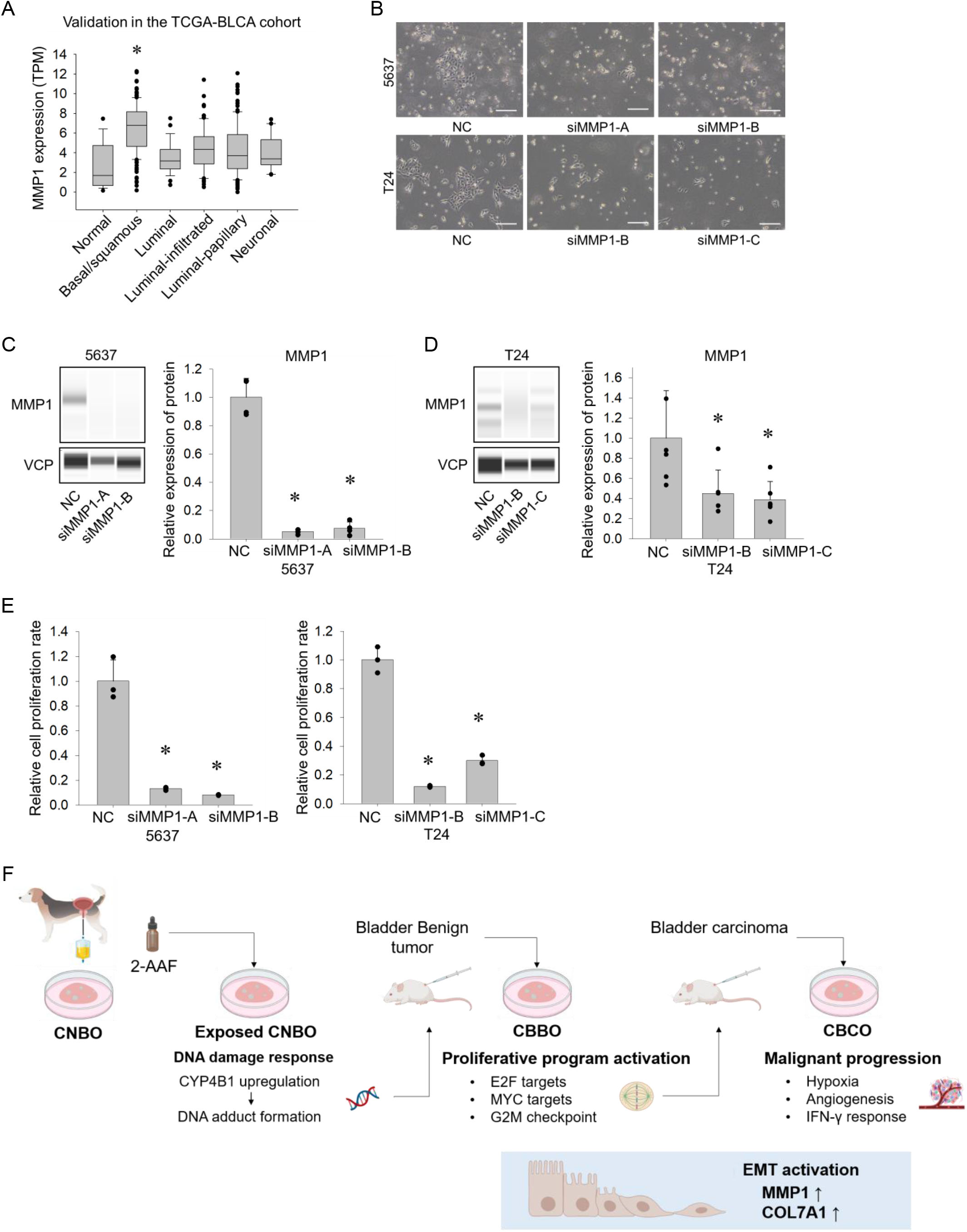
Validation of MMP1 function in human bladder cancer cells. (A) MMP1 expression in normal bladder tissue and molecular subtypes of bladder cancer in the Cancer Genome Atlas-Bladder Urothelial Carcinoma (TCGA-BLCA) cohort. (B) Representative bright-field images of 5637 or T24 cells transfected with NC or two independent siRNAs targeting *MMP1*. Scale bar, 300 µm. (C, D) Western blot analysis and quantification of MMP1 expression in 5637 and T24 cells following transfection with control or MMP1-targeting siRNAs. VCP was used as a loading control. Data are presented as mean ± SD (n = 4 for 5637 cells and n = 6 for T24 cells). (E) Quantification of cell proliferation in 5637 and T24 cells transfected with NC or siRNAs targeting MMP1. Data are presented as mean ± SD and relative values normalized to NC (n = 3). (F) Schematic overview of the stepwise bladder tumorigenesis model established in this study. The diagram summarizes the progression from CNBO to CBBO and CBCO and the major molecular and phenotypic changes associated with tumor progression, including EMT activation and the functional involvement of MMP1 and COL7A1.

## Discussion

In this study, we reconstructed stepwise bladder carcinogenesis from normal urothelium through benign tumor formation to invasive carcinoma and longitudinally characterized the molecular changes accompanying malignant progression. A key finding was that tumor progression occurred despite relatively limited acquisition of canonical cancer-associated genomic alterations and was instead accompanied by extensive transcriptional reprogramming, including enrichment of EMT-associated programs. Functional analyses identified COL7A1 and MMP1 as mediators of tumor cell proliferation and invasion, and these dependencies were further validated in independently established organoids from spontaneous canine bladder cancers. Notably, MMP1 knockdown also reduced COL7A1 expression, suggesting a potential regulatory relationship between these molecules. The translational relevance of these findings was further supported by the association of MMP1 with basal/squamous human bladder cancer and by its functional contribution to the proliferation of human bladder cancer cells. Together, these findings highlight extensive transcriptional reprogramming, rather than the progressive accumulation of canonical genomic alterations alone, as a prominent feature of malignant progression and identify MMP1 and COL7A1 as functional mediators of malignant phenotypes.

Although metabolic activation is generally required for 2-AAF carcinogenicity, aromatic amine carcinogens can be metabolized not only in the liver but also in bladder tissues^75^. In CNBO, 2-AAF exposure induced expression of *CYP4B1*, *HMOX1*, and *NQO1*, together with activation of pathways related to xenobiotic metabolism, glutathione metabolism, and detoxification (Fig. 1E-G). Furthermore, AAF-DNA adduct formation and increased γH2AX positivity were observed following continued exposure (Fig. 1H, I). These findings suggest that CNBO can mount molecular responses to aromatic amine exposure that are accompanied by DNA damage. Importantly, these responses were observed in organoids cultured in the absence of exogenous metabolic activation systems, indicating that bladder organoids can be used to investigate early cellular responses to aromatic amine carcinogens.

Comparison of CNBO, CBBO, and CBCO revealed progressive phenotypic changes during tumor progression. CNBO exhibited the highest organoid growth, whereas growth was reduced in both CBBO and CBCO (Fig. 2I, J). Organoids are self-organizing three-dimensional epithelial structures that recapitulate key architectural features of epithelial tissues^60^. Therefore, the reduced organoid growth observed in CBBO and CBCO may reflect alterations in epithelial organization accompanying tumor progression^76,77^. Consistent with these phenotypic changes, differences in drug sensitivity were also observed, particularly for vinblastine and lapatinib. Together, these findings support the notion that CNBO, CBBO, and CBCO represent biologically distinct cellular states during stepwise tumor progression.

Whole-exome sequencing showed that BRAF mutations, which are frequently observed in canine urothelial carcinoma^17–20^, were not detected in CBCO. Furthermore, no clear mutations were identified in Tier 1 bladder cancer–related genes in the Cancer Gene Census (CGC)^78^, and the tumor mutational burden (TMB) remained relatively low. These findings suggest that tumor progression in this model was not primarily driven by the accumulation of canonical bladder cancer driver mutations.

In contrast to the relatively stable genomic landscape, transcriptomic analysis revealed marked changes during tumor progression. EMT-related gene sets showed a stepwise increase from CNBO to CBBO and CBCO, suggesting that malignant progression in this model is characterized by extensive transcriptional reprogramming despite relatively limited acquisition of canonical cancer-associated genomic alterations. EMT has been implicated not only in invasion and metastasis but also in early stages of tumor development and cellular plasticity^79^. Consistent with this, *COL7A1* and *MMP1* were progressively upregulated during tumor progression and were functionally associated with cell proliferation and invasion. Stage-specific analyses further indicated that the transition from CNBO to CBBO was characterized primarily by enrichment of cell cycle–related programs, including E2F targets, MYC targets, and G2M checkpoint (Supplementary Fig. S2A), whereas progression from CBBO to CBCO was associated with gene sets related to adaptation to the tumor microenvironment, including hypoxia, angiogenesis, and inflammatory signaling pathways (Supplementary Fig. S2B). These findings suggest that tumor progression in this model involves sequential acquisition of proliferative and adaptive characteristics accompanied by EMT activation. Taken together, these findings support a model in which bladder tumor progression is accompanied by sequential transcriptional reprogramming, EMT activation, and upregulation of COL7A1 and MMP1.

To determine whether the molecular features identified in the stepwise carcinogenesis model are shared with naturally occurring disease, we analyzed organoids derived from spontaneous canine bladder tumors (Fig. 6). Knockdown of either *COL7A1* or *MMP1* reduced cell proliferation and tumorigenic capacity, supporting their contribution to tumor growth (Fig. 6D, F). RNA-seq analysis further demonstrated downregulation of mitosis-related gene sets following knockdown of either gene, suggesting a role for these genes in maintaining proliferative activity. Interestingly, distinct transcriptional responses were observed following knockdown of each gene. *COL7A1* knockdown was associated with enrichment of extracellular matrix– and tissue organization–related pathways (Supplementary Fig. S3A), whereas *MMP1* knockdown preferentially enriched innate immune and interferon-related programs (Supplementary Fig. S3B). These findings suggest that, although both genes contribute to tumor growth, they may influence tumor progression through partially distinct molecular mechanisms. Notably, MMP1 knockdown also reduced COL7A1 expression, suggesting a potential regulatory relationship between these molecules. Given the established roles of MMP1 in extracellular matrix degradation and tumor-associated tissue remodeling and of COL7A1 as a structural component of the extracellular matrix, further studies will be required to determine whether and how MMP1 regulates COL7A1 during bladder cancer progression^80^.

Analyses of CBCO and sCBCO identified *MMP1* as a molecule upregulated during bladder cancer progression (Fig. 5, 6). Consistent with these findings, TCGA analysis demonstrated elevated *MMP1* expression in the basal/squamous subtype of human bladder cancer (Fig. 7A). Interestingly, the tumors from which CBCO was established exhibited histological evidence of squamous differentiation. In addition, RNA-seq analysis of CBCO revealed increased expression of several genes associated with basal and squamous phenotypes, including *KRT5*, *KRT6A*, *KRT14*, and *EGFR*. Together, these findings suggest that CBCO shares certain molecular and histopathological characteristics with the basal/squamous subtype of human bladder cancer. Consistent with our findings, *MMP1* expression has been reported to increase with bladder cancer progression and to be associated with tumor invasion^81^. Functional validation in human bladder cancer cell lines further demonstrated that MMP1 knockdown reduced cell proliferation, supporting the cross-species relevance of MMP1 in bladder cancer. However, the clinical significance and mechanistic role of MMP1 in human bladder cancer require further investigation.

This study has several limitations. First, the stepwise model was established from a single normal canine bladder organoid line derived from one healthy dog, and serial xenotransplantation may have imposed selective pressures that contributed to the phenotypic and molecular changes observed during progression. Thus, the changes identified in CBBO and CBCO cannot be attributed exclusively to 2-AAF-induced carcinogenesis. Nevertheless, the functional relevance of COL7A1 and MMP1 was reproduced in independently established organoids from spontaneous canine bladder cancers, supporting the broader relevance of these molecular alterations. Future studies using additional normal organoid lines and independently generated stepwise models will be required to determine the generalizability of the molecular trajectories identified here.

In conclusion, by reconstructing stepwise bladder carcinogenesis from normal urothelium to invasive carcinoma, we identified extensive transcriptional reprogramming, including EMT activation, during malignant progression despite relatively limited acquisition of canonical cancer-associated genomic alterations. Functional analyses further identified COL7A1 and MMP1 as mediators of malignant phenotypes, with these dependencies validated in independently established spontaneous canine bladder cancer organoids. The relevance of MMP1 was further supported in human bladder cancer, where it was associated with the basal/squamous subtype and contributed to tumor cell proliferation. Together, these findings highlight transcriptional reprogramming as a prominent feature of bladder cancer progression and identify MMP1 and COL7A1 as functional mediators of malignant phenotypes.

## Acknowledgements

This work was supported by a Grant-in-Aid for Scientific Research (A) from the Japan Society for the Promotion of Science (JSPS KAKENHI Grant Number JP24H00540) to T.U.

## Data availability

The raw RNA-sequencing and whole-exome sequencing data generated in this study have been deposited in the NCBI Sequence Read Archive (SRA). The four datasets generated in this study are scheduled to be consolidated under BioProject accession number PRJNA1519075. The RNA-sequencing datasets include data from 2-AAF-treated CNBO, CNBO, CBBO, CBCO, and siRNA-treated spontaneous canine bladder cancer organoids. The whole-exome sequencing datasets include CNBO, CBBO, and CBCO. All other data supporting the findings of this study are available within the article and its Supplementary Information or from the corresponding author upon reasonable request.

